# Mis-annotated multi nucleotide variants in public cancer genomics datasets can lead to inaccurate mutation calls with significant implications

**DOI:** 10.1101/2020.06.05.136549

**Authors:** Sujaya Srinivasan, Natallia Kalinava, Rafael Aldana, Zhipan Li, Sjoerd van Hagen, Sander Y.A. Rodenburg, Megan Wind-Rotolo, Ariella S. Sasson, Hao Tang, Xiaozhong Qian, Stefan Kirov

## Abstract

**Background:** Next generation sequencing is widely used in cancer to profile tumors and detect variants. Most somatic variant callers used in these pipelines identify variants at the lowest possible granularity – single nucleotide variants (SNVs). As a result, multiple adjacent SNVs are called individually instead of as a multi-nucleotide variant (MNV). The problem with this level of granularity is that the amino acid change from the individual SNVs within a codon could be different from the amino acid change based on the MNV that results from combining the SNVs. Most variant annotation tools do not account for this, leading to incorrect conclusions about the downstream effects of the variants.

**Method:** Here, we used Variant Call Files (VCFs) from the TCGA Mutect2 caller, and developed a solution to merge SNVs to MNVs. Our custom script takes the phasing information from the SNV VCFs and based on a gene model, determines if SNVs are at the same codon and need to be merged into a MNV prior to variant annotation.

**Results:** We analyzed 10,383 VCFs from TCGA and found 12,141 MNVs that were incorrectly annotated. Strikingly, the analysis of seven commonly mutated genes from 178 studies from cBioPortal revealed that MNVs were consistently missed in 20 of these studies, while they were correctly annotated in 15 more recent studies. The best and most common example of MNVs was found at the BRAF V600 locus, where several public datasets reported separate BRAF V600E and BRAF V600M variants, instead of a single merged V600K variant.

**Conclusion:** While some datasets merged MNVs correctly, many public datasets have not been corrected for this problem. As a best practice for variant calling, we recommend that MNVs be accounted for in NGS processing pipelines, thus improving analyses on the impact of somatic variants in cancer genomics.

## Background

Next generation sequencing is commonly used in cancer to determine the underlying genomic features of the tumor^1^. Pipelines that convert the raw sequencing data into useful knowledge include sequence alignment, variant calling and annotation tools. Single nucleotide variants and indels are the most common type of variants called by most variant callers, and these variants are prevalent in many important cancer genes. Most popular variant callers like Mutect2^2^, VarScan2^3^, VarDict^4^, strelka2^5^ and the Sentieon^6^ suite of tools call variants at the most granular level of single nucleotide variants (SNVs) and indels. Missense and nonsense variants produce amino acid changes that could result in a protein that is either non-functional, or has a different or impaired function. Accurate annotation, of the amino acid changes that occur due to the SNVs and indels, is therefore critical to understanding the functional consequences of these variants.

A multi-nucleotide variant (MNV) is defined as two or more variants within the same codon on the same haplotype (see Figure 1). Variant callers commonly detect SNVs and small indels, but most callers and downstream variant annotation tools fail to consider whether nearby variants are part of the same haplotype. If multiple nearby variants happen to be within a single codon, the amino acid change could be different from the individual amino acid changes resulting from the SNVs. Many variant callers, such as Strelka, VarScan and VarDict, do not include haplotype or phase information with the variant calls. Some of the more recent variant callers such as Mutect2, Sentieon TNScope and Sentieon TNHaplotyper include phase information to indicate if nearby variants are in phase (i.e. part of the same haplotype) when there is enough evidence from the reads supporting the variants.

**Figure 1:**
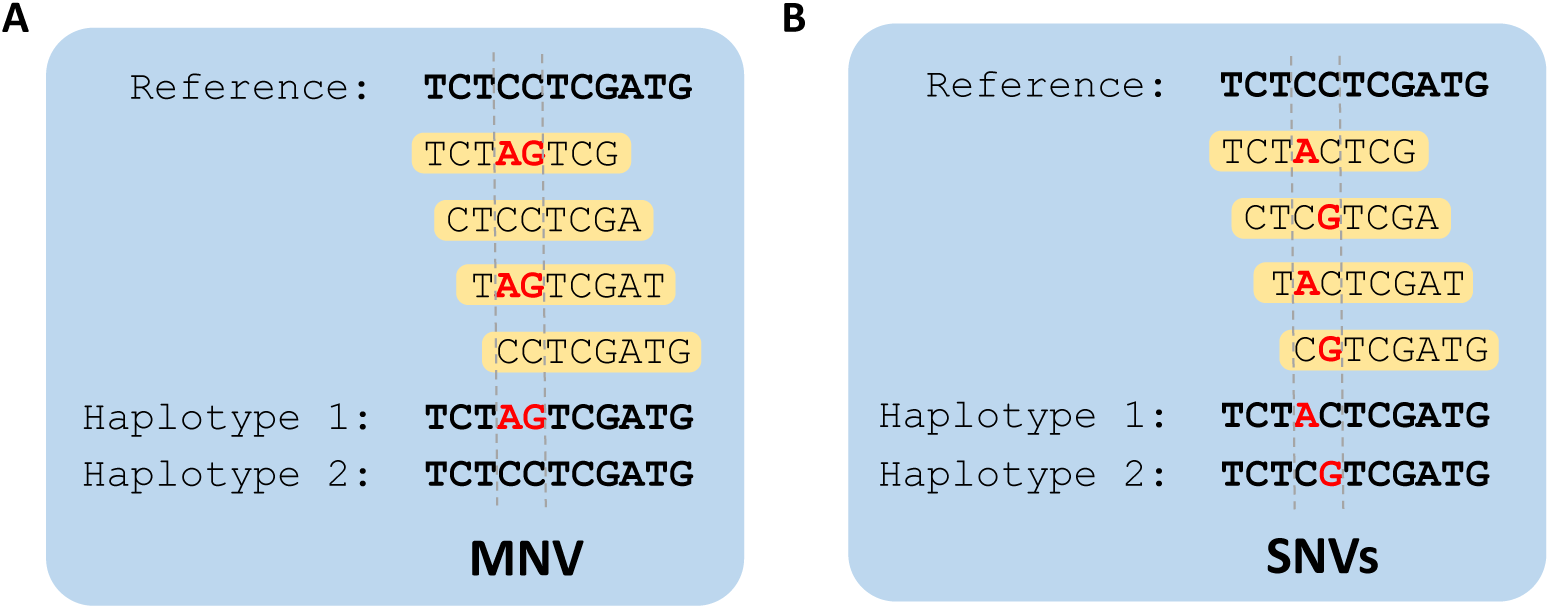
Schematic presentation of MNV and SNV events. (A) The 2 SNVs co-occurring on the same reads indicates they are part of the same haplotype and should be annotated as MNV. (B) The 2 SNVs in this case are adjacent but on different reads, and should be annotated as individual SNVs.

Commonly used variant annotation tools, such as SnpEff^7^, ANNOVAR^8^, & Ensembl Variant Effect Predictor (VEP)^9^, annotate variants individually without considering haplotype information or combining nearby in-phase variants to MNVs. There are some tools such as bcftools csq^10^ (haplotype aware consequence caller) that have tried to address this problem, but the software expects phased VCFs as input with phasing information in the genotype (GT) field in a specific and seldom used format. MAC ^11^(Multi-nucleotide Variant Annotation Corrector) requires both the VCF and the corresponding Binary Alignment Map (BAM) file in order to correct for MNVs, corresponding to adjacent SNVs. MACARON (Multi-bAse Codon Association variant ReannotatiON)^12^ is another tool that uses both the VCF and the BAM to re-annotate VCFs with corrected MNVs from multiple SNVs within a codon.

There are several important cancer genes that are known to have hotspot regions with many variants. A few examples are BRAF at the V600 locus, and KRAS at G12 and G13 loci. Sometimes these variants are part of the same haplotype, and therefore should be annotated as MNVs, but most pipelines annotate them as multiple SNVs. This could lead to incorrect functional predictions for the effect of the variants.

In this paper, we consider some common public cancer genomics datasets to understand if MNVs are accounted for, and propose a method to merge SNVs into MNVs.

## Results

### TCGA results

We downloaded 10,383 Mutect2 VCF files processed with the human reference genome (GRCh38) from The Cancer Genome Atlas (TCGA). The downloaded VCFs comprise 33 cancer types or indications.

We post-processed the TCGA mutect2 VCFs using a custom developed MNV merge script. This script takes the SNVs that are in phase and within the same codon and merges them into MNV. We excluded repeat regions and major histocompatibility complex (MHC) regions for this analysis, and only characterized the instances of merged SNVs. Indels were not considered at this time. We found that across all files, there were a total of 12,141 MNVs that were originally annotated as multiple SNVs, and of these 6,357 had a completely novel protein effect, i.e the new protein effect was different from the SNVs’ protein effects (Table 1, Fig. 2). The most frequent novel MNV events were new missense events (5,413). Nonsense events, both stop gain (254) and rescue of nonsense (517), had the most impact on the interpretation of protein function. This shows that annotating MNVs correctly can significantly alter downstream analysis results.

**Table 1:** Most commonly mis-annotated MNVs in cBioPortal among the 7 genes that were studied.

| Gene | SNVs | MNV | Count |
| --- | --- | --- | --- |
| BRAF | V600M + V600E | V600K | 52 |
| BRAF | V600M + V600G | V600R | 9 |
| BRAF | G469V + G469* | G469L | 2 |
| KRAS | G12V + G12C | G12F | 8 |
| KRAS | G12A + G12C | G12S | 2 |
| KRAS | G12V + G12S | G12I | 2 |
| KRAS | G12V + G12R | G12L | 2 |
| NRAS | Q61R + Q61K | Q61R | 5 |

**Figure 2:**
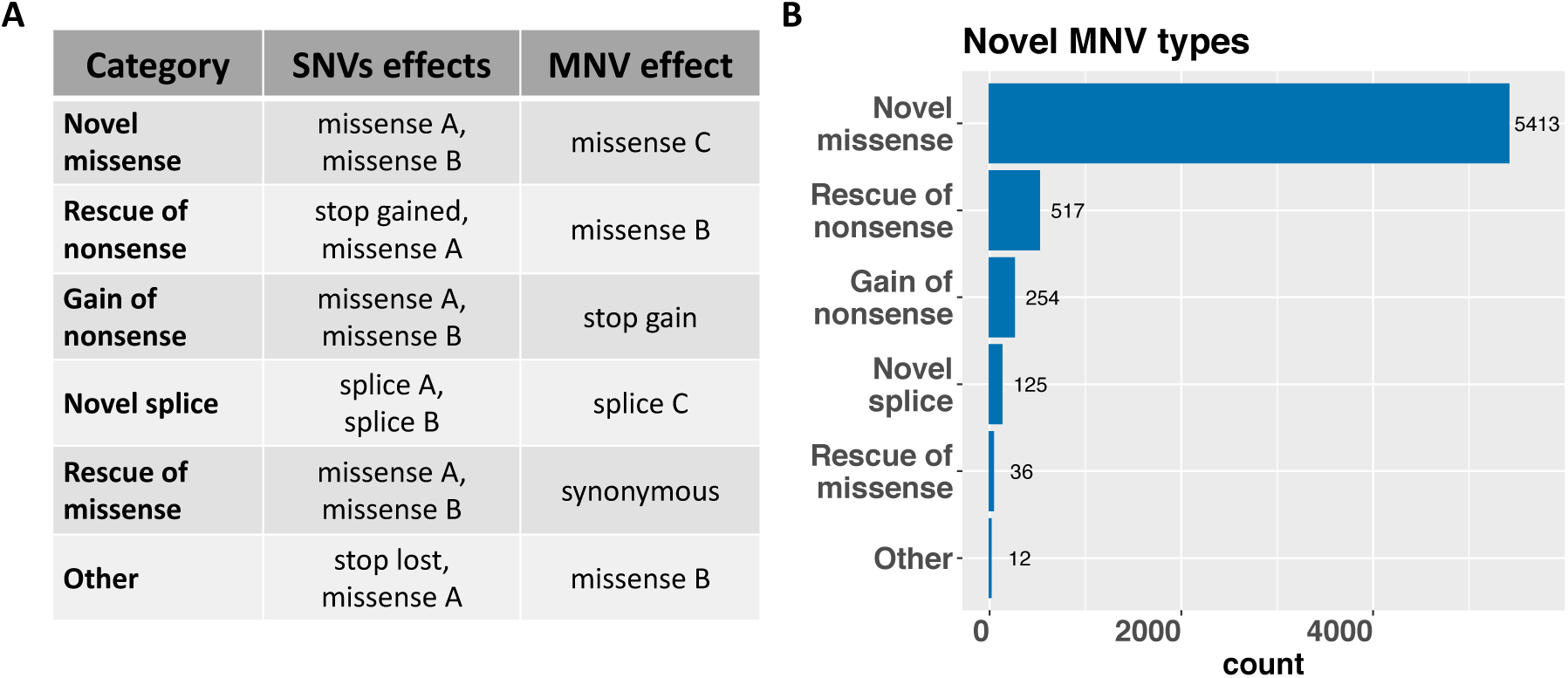
Novel MNV effects in TCGA data. (A) Categories and examples of the MNV novel annotation effect as a result of combination of two SNVs. (B) Number of MNVs for novel effects in TCGA data.

Skin Cutaneous Melanoma (SKCM) and lung cancers: lung adenocarcinoma (LUAD) and lung squamous cell carcinoma (LUSC) had the highest percentage of samples with MNVs (Fig. 3a). We also found the highest median number of SNVs and MNVs in SKCM, LUAD and LUSC (Fig. 3b and c). This is expected because of the high Tumor Mutation Burden (TMB) in these indications. Breast cancer (BRCA), the indication with the largest number of samples in this dataset (1,040 samples), is known to have a low TMB^13^, and our results are consistent with this.

**Figure 3:**
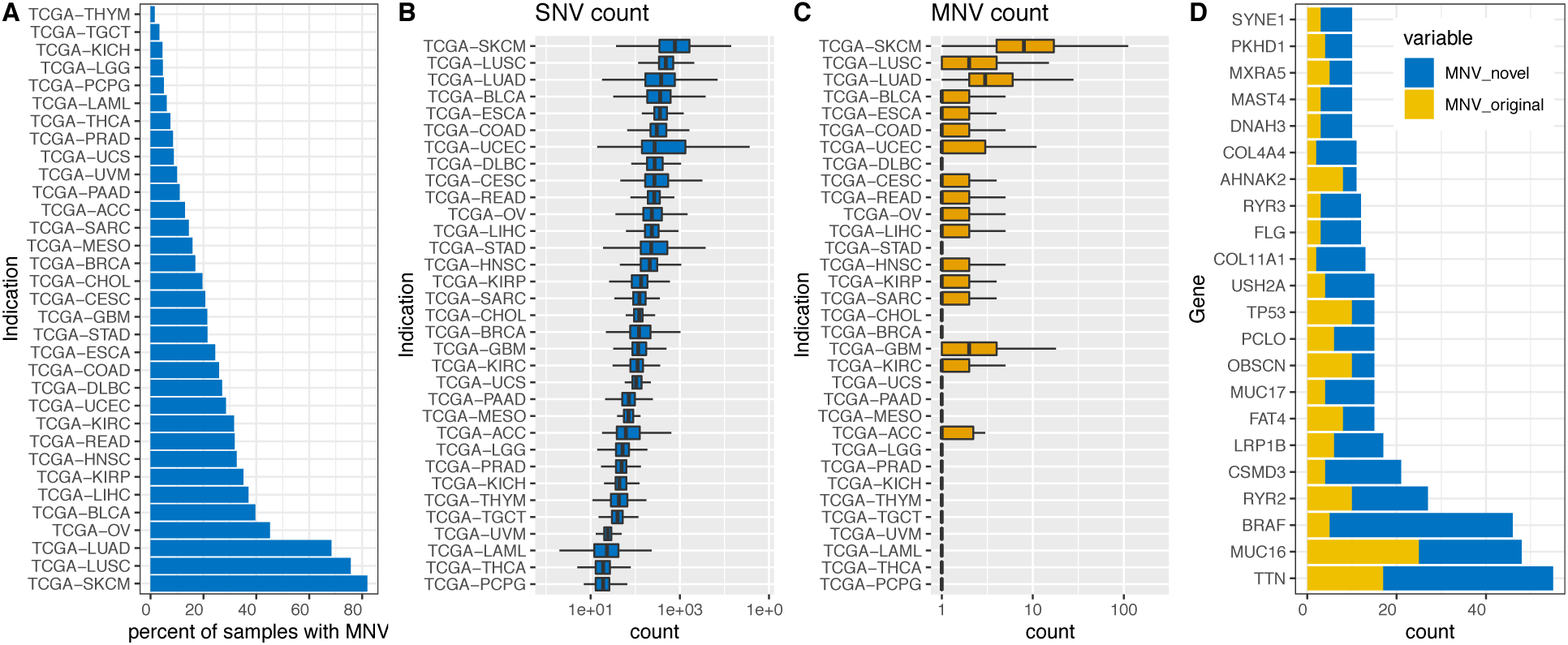
MNV summary in TCGA dataset. (A) Distribution of TCGA samples by indication. The bars indicate the percent of samples that had MNV(s). (B) Boxplot of the SNV count per indication. (C) Boxplot of the MNV count per indication. Indications are ordered the same as the SNV count. (D) Distribution of novel and original MNV for genes with total MNV ≥ 10.

While most genes had only one or two MNVs, we found 22 genes that had 10 or more MNVs (Fig. 3d). Many of these genes are known for hotspot mutations, so this finding is not that surprising. The most consistent MNVs were in the BRAF gene: 43 out of 46 MNVs were at the V600 locus, all with a novel missense outcome. Furthermore, a single BRAF V600M never occurred alone, but always co-occurred in phase with another variant V600G or V600E, leading to the novel mutations V600R and V600K respectively.

### cBioPortal results

We analyzed mutation annotation files (MAF) from cBioPortal^14,15^ (http://www.cbioportal.org) from all non-redundant studies (178) for 7 cancer genes (BRAF, KRAS, NRAS, PTEN, BRCA1, BRCA2, MUC16). Since cBioPortal MAFs do not have phasing information, we used counts for the variant reads and variant allele frequencies, as proxies for phase. If the variant allele frequencies of two variants within a codon was approximately the same, we inferred that they co-occurred on the same read (Fig. 4).

**Figure 4:**
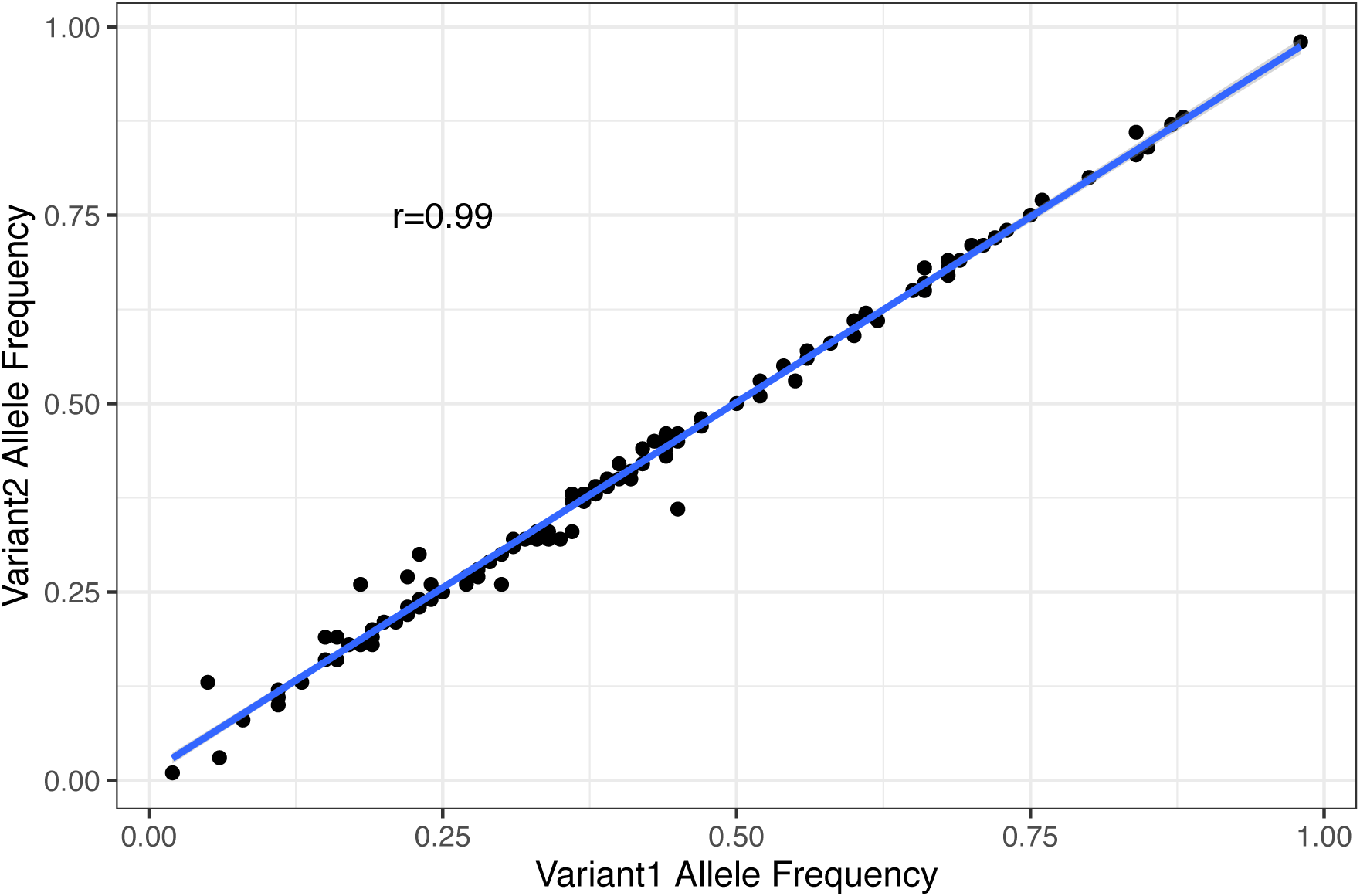
Variant allele frequencies of variants present on the same codon in cBioPortal. The high correlation between the VAFs of the variants indicates that they were present on the same reads.

Some common hotspot regions of cancer genes, like BRAF V600 and KRAS G12 loci, were particularly affected by not merging the SNVs into MNVs. While some studies did call the MNVs correctly, there were 20 studies, including several TCGA studies, that did not. Table 1 shows the most common mis-annotated MNVs among the seven genes that we studied. The most frequently mis-annotated MNV was at the BRAF V600 locus, with a total of 61 MNVs (V600K and V600R). The KRAS G12 locus had 14 MNVs with the most common being co-occurring G12V and G12C SNVs which should have been annotated as G12F.

In our analysis of all BRAF V600 variants from cBioPortal, we found only two occurrences of V600M alone, with no other variant. Since we did not have the full set of variant calls from this dataset, it was not possible for us to determine if these two occurrences were actually V600M, or if they co-occurred with another variant that was filtered out for quality reasons, or due to the fact that it was a synonymous variant. There were 64 other samples that had a BRAF V600M variant, but those samples also had either a V600G or V600E variant (Supplementary Table 1). When we examined all studies, including duplicate samples from studies that were submitted at different times, we found that there were conflicting entries for some samples. The SNVs from the earlier submissions were replaced by MNVs in later submissions, indicating that pipelines had probably been updated to correct for MNVs. Some examples of these are the corrections for the KRAS G12 variants and the BRAF V600 variants (Supplementary table 1). One important example of a corrected MNV was a V600D, which consists of a synonymous variant along with a V600E. These variants would be completely missed in our analysis from cBioPortal, since synonymous variants are filtered out. They would only appear if MNVs were correctly handled.

### Double base mutation patterns

Somatic variants in cancer genomes have specific patterns, known as Mutational Signatures^16^ associated with underlying processes that characterize the specific etiology of the cancer. The Doublet Base Substitution (DBS) Signatures published in Mutational Signatures v3^17^ show the two base-pair signatures that are characteristic of certain cancer types.

We analyzed the TCGA data after it had been corrected for MNVs, and identified the most common double-base mutation patterns in Fig. 5a. We found that the CC to TT change was prominent in melanoma samples (Fig. 5b). This is consistent with the reported signature, DBS 1, which is a characteristic of UV related damage. Lung cancer (LUAD and LUSC) samples predominantly showed CC to AA change (Fig. 5b), which was consistent with the DBS 2 signature, indicating exposure to tobacco smoking^17^. This shows that the detected MNVs are consistent with the expected mutational signatures, and by analyzing MNVs, we can detect underlying patterns that would be missed otherwise.

**Figure 5:**
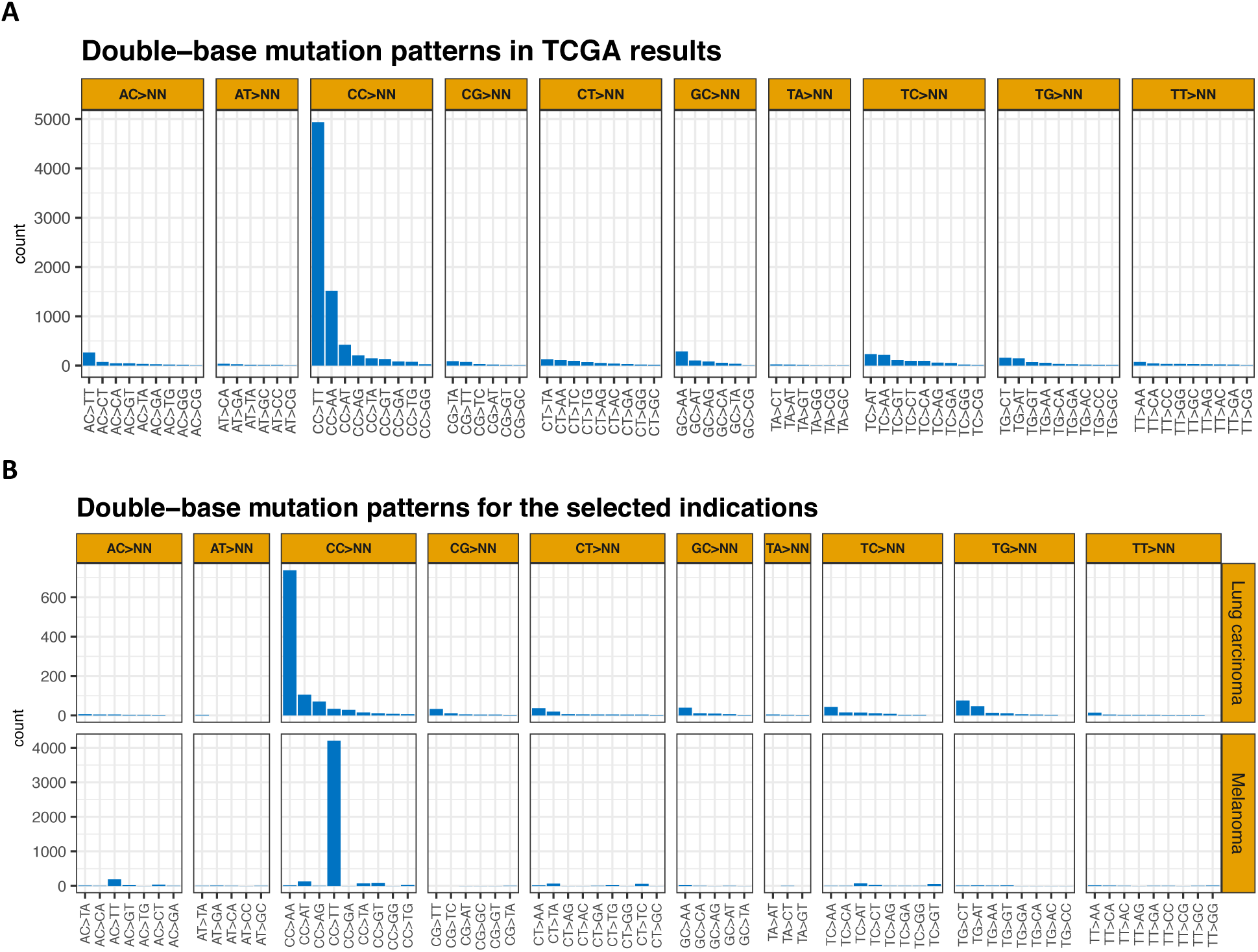
Double-base mutation patterns found in the TCGA data based on the MNV corrections. (A) Frequency of double-base mutation patterns found in all indications of TCGA results. The reverse complement was accounted according to the double-base signatures described in Alexandrov et al, 2020. (B) Double-base mutation patterns plotted for the selected indications: melanoma and lung carcinoma. Lung adenocarcinoma (LUAD) and lung squamous cell carcinoma (LUSC) were combined into one panel for lung carcinoma.

## Discussion

We analyzed VCFs from TCGA as well as MAF files from cBioPortal, and found that there were over 12,000 MNVs that were characterized as SNVs in TCGA. Many of these MNVs are in important cancer genes, such as BRAF and KRAS. From a functional perspective, it is important to annotate these variants correctly, so that the effects of the variants can be properly evaluated and interpreted. For example, we did not find a single occurrence of a BRAF V600M alone in any of the studies, it was always in phase with a V600E or V600G.

At the same time a number of publications reference V600M^18–28^; COSMIC database at the time we reviewed the data lists 31 occurrences of V600M. The methods for detecting the mutation are extremely diverse, ranging from Restriction Fragment Length Polymorphism(RFLP) and direct Sanger sequencing to MassArray/Sequenom platform. We cannot evaluate to what extent these methods have the ability to detect MNVs as this is beyond the scope of this study, but it likely these errors are more broadly occurring.

Other studies identified double V600M-V600E or V600M-V600G mutants that are possibly MNVs, as the detection method does not allow for phasing information to be known (typically Sanger sequencing)^29,30^. In this specific case, the correct identification of the amino acid change may have serious consequences. A number of BRAF inhibitors are approved for either V600E or V600E/K^31,32^, but treatment options may differ for other rare mutations, including V600M. For example, there is preclinical data suggesting that BRAF kinase activity may not be altered in V600M/A unlike V600E/K/D^33^. Retrospective analysis points to V600K carriers having a worse prognosis^34^ and worse PFS response to existing BRAF inhibitors^35^.

From a cancer biology perspective, it is also curious to understand how these MNVs evolve. The V600K/R for example are not driven by UV damage as V600K originates from GT->AA and V600R originates from GT->AG, whereas UV signature is associated with C->T events^36^. Studies on germ-line MNVs have shown that this type of events tends to be more pathological than SNVs and associated mostly with APOBEC and DNA polymerase zeta^37^. Another potential mechanism would argue that two independent SNVs happen to occur by chance in the same codon, and that the resulting MNV clone gains an advantage and eventually displaces the original SNV clone from the tumor population. However, we should be able to at least occasionally detect the founding clone mutation in the same tumor specimen, evidence of which we have not seen to date.

There have been many large-scale efforts to characterize MNVs within a germline context, most recently with gnomAD^38^. However, one of the potential issues we did not address in this paper is when germline variants are part of the same haplotype with a proximal somatic variant as part of the same codon. There is not much evidence that this is a widespread problem^39^, but it would be important to assess the effect of it.

In our analysis of the various cBioPortal studies, we observed that several studies after 2017 from larger academic hospitals and institutions had corrected for MNVs, indicating that the problem was recognized and fixed in some of these pipelines. We also found that the ICGC PCAWG^40^ effort and the AACR Genie^41^ project called MNVs correctly. However, there are still several smaller academic and commercial labs that may not have fixed this issue, and our analysis shows the need for the MNV merge step to be incorporated into variant-calling pipelines as a standard best practice. Needless to say, clinical assays should be assessed not only on the correct characterization of BRAF V600 mutants, but also the precise amino acid change associated with it.

## Methods

### MNV Merging for TCGA VCFs

We downloaded 10,383 TCGA VCFs processed using the Mutect2 variant caller on the GRCh38 reference genome from the Cancer Genomics cloud. When nearby variants are part of the same haplotype (in phase), Mutect2 adds tags to indicate this – PGT is the phased genotype of the variant, and PID is an ID that is shared between variants of the same haplotype; this information is then used by a python script to merge SNVs to MNVs.

We downloaded Refseq transcripts BED file from the UCSC table browser (https://genome.ucsc.edu) and pre-processed it into a codon file that had the positions of each codon defined. The MNV merge script then used this codon file to determine whether to merge SNVs, based on whether they are part of the same haplotype and codon.

The python script (merge_mnp.py) takes the input VCF, reference genome, pre-processed codons text file and a parameter that specifies if indels should be considered. For the purposes of this study, we did not consider indels. The python script identifies SNVs that are both in phase and within the same codon into a new MNV. The new MNV has a PASS in the filter field, while the original SNVs have a MERGED in the filter field to represent that they have been superseded by the MNV. All code can be found on GitHub at https://github.com/Sentieon/sentieon-scripts.

The VCFs that have the merged MNVs were annotated using SnpEff. Annotations from gnomAD v2.1.1 ^42^, dbSNP^43^ version 146 and COSMIC^44^ version 84 were added to the VCFs, and both “PASS” and “MERGED” variants were retained in order to be able to trace the MNVs and the original SNVs. The repeat masker GRCh38 annotations were used to mask the repetitive regions and were excluded from the MNV analysis. The highly variable MHC region at chromosome 6 position 28510120 – 33480578 was also excluded from the MNV analysis.

### cBioPortal

We downloaded Mutation Annotation Files (MAF) from the cBioPortal (https://www.cbioportal.org/) by choosing “Curated list of non-redundant studies” for 7 genes – BRAF, KRAS, NRAS, PTEN, BRCA1, BRCA2, MUC16. To identify variants that were part of the same haplotype and at the same codon position, we looked for those instances where there were multiple variants from the same sample at the same codon position, and had the same Variant Allele Frequency (VAF). This indicated that it was highly likely that the variants appeared together on most reads

In addition, we queried the public cBioPortal API (https://www.cbioportal.org/api/), retrieving the complete collection of mutation data for all loaded studies. We then filtered mutations for few selected mutation hotspots, i.e. BRAF V600, KRAS G12, and NRAS Q61, and subsequently determined which variant calls occurred in each sample at these hotspots. Samples occurring in multiple studies were combined, but we kept track of the cases where samples had different variant calls between studies.

## Declarations

### Ethics approval and consent to participate

Not applicable

### Consent for Publication

Not applicable

### Availability of data and materials

TCGA data is available from the Genomic Data commons at https://portal.gdc.cancer.gov/. Data from cBioPortal is available at https://www.cbioportal.org/.

### Financial & competing interests disclosure

The research for this paper was funded by Bristol Myers Squibb. Rafael Aldana and Zhipan Li are employees of Sentieon, Inc. Sjoerd van Hagen and Sander Y.A. Rodenburg are employees of The Hyve. Xiaozhong Qian is an employee of Daichi Sankyo, Inc. The other authors have no conflicts of interest to declare. No writing assistance was utilized in the production of this manuscript.

### Author contributions

SS and NK identified, processed and analyzed the datasets, and wrote the manuscript. SK identified the problem presented in the manuscript and SK, SS and NK conceived the idea. RA and ZL developed the scripts and code. SH and SR helped with data retrieval and analysis from cBioPortal. MWR, AS, HT and XQ contributed to the analysis of the results and downstream implications. All authors discussed results and contributed to the final manuscript.

## Acknowledgements

The results shown here are in whole or part based upon data generated by the TCGA Research Network: https://www.cancer.gov/tcga.

## Ethical conduct of research

The authors state that they have obtained appropriate institutional review board approval or have followed the principles outlined in the Declaration of Helsinki for all human or animal experimental investigations. In addition, for investigations involving human subjects, informed consent has been obtained from the participants involved.

---

**Supplementary Table 1** Table of all samples from cBioPortal that have a variant at the BRAF V600 and G469, KRAS G12 and NRAF Q61 loci. The common samples that have conflicting annotations between studies are indicated by separating with a “;”

## Notes

https://portal.gdc.cancer.gov/

https://www.cbioportal.org/

### Summary of Updates

Author affiliation corrected.

